# Understanding the Immunomodulatory Effects of Bovine Colostrum: Insights into IL-6/IL-10 Axis-Mediated Inflammatory Control

**DOI:** 10.1101/2023.07.26.550707

**Authors:** Ramune Grigaleviciute, Paulius Matusevicius, Rita Planciuniene, Rolandas Stankevicius, Eivina Radzeviciute-Valciuke, Austeja Baleviciute, Augustinas Zelvys, Aukse Zinkeviciene, Vilma Zigmantaite, Audrius Kucinskas, Povilas Kavaliauskas

**Affiliations:** Biological Research Center, Lithuanian University of Health Sciences, Tilzes str. 18/7, Kaunas, LT-47181, Lithuania; Department of Animal Nutrition, Lithuanian University of Health Sciences, Tilzes str. 18 Kaunas, LT-47181, Lithuania; Institute of Microbiology and Virology, Lithuanian University of Health Sciences. Eiveniu str. 4, Kaunas, LT-50161, Lithuania; Centre for Innovative Medicine, department of immunology, Santariskiu str. 5, Vilnius, LT-08410, Lithuania; Institute of Environmental Medicine, Toxicology Unit, Karolinska Institutet, Stockholm, Solnavägen 1, 17177, Solna, Sweden; Joan and Sanford I. Weill Department of Medicine, Weill Cornell University, 1300 York Avenue, NY 1109. New York, United States; Department of Microbiology and Immunology, University of Maryland Baltimore School of Medicine, Baltimore, MD 21201, USA; Institute of Infectious Diseases and Pathogenic Microbiology, Birstono str. 38A, LT-59116, Prienai, Lithuania

**Keywords:** bovine colostrum, cytokine production, immunomodulation, intestinal permeability, THP-1 macrophages, Caco-2 cells, NF-κB, *S. typhimurium* infection, inflammatory disorders

## Abstract

Bovine colostrum (COL), the first milk secreted by lactating cows postpartum, is a rich source of bioactive compounds that exert significant role on the survival, growth, and immune development of neonatal calves [9,10]. This study investigated the immunomodulatory effects of COL on cytokine production *in vitro* using a Caco-2/THP-1 macrophage co-culture model stimulated with Phorbol 12-myristate 13-acetate (PMA). COL pretreatment significantly reduced IL-6 production induced by PMA, while increasing IL-10 production. Further investigations revealed that the IL-6 suppressive effect of colostrum was heat-sensitive and associated with components of higher molecular mass (100 kDa). Moreover, colostrum primarily influenced THP-1 macrophages rather than Caco-2 epithelial cells. The effects of colostrum on IL-6 production were associated with reduced NF-κB activation in THP-1 macrophages. In calf-FMT transplanted C57BL/6 murine model, colostrum decreased intestinal permeability, reduced immune cell infiltration, and suppressed IL-6 production during *S. typhimurium* infection. These results highlight the immunomodulatory activity of bovine colostrum and its potential therapeutic applications in inflammatory disorders. Further studies are needed to elucidate the underlying mechanisms and validate the findings in bovine models.

**Simple Summary:** This study explores the immunomodulatory properties of bovine colostrum (COL), the initial milk produced by lactating cows, on cytokine production *in vitro* and in a novel murine calf-FMT model. The researchers utilized a Caco-2/THP-1 macrophage co-culture model stimulated with Phorbol 12-myristate 13-acetate (PMA) to investigate the effects of COL on cytokine production. The findings indicate that COL pretreatment significantly reduced IL-6 production while enhancing IL-10 production. The IL-6 suppressive effect was heat-sensitive and associated with components of higher molecular mass (100 kDa). Colostrum demonstrated decreased intestinal permeability, reduced immune cell infiltration, and suppressed IL-6 production during *S. typhimurium* infection. These results highlight the immunomodulatory potential of bovine colostrum and its prospective therapeutic applications in inflammatory disorders. Further research is necessary to elucidate the underlying mechanisms and corroborate the findings in bovine models.

## 1. Introduction

Bovine colostrum (COL), the first milk produced by lactating cows postpartum, is a rich source of bioactive compounds that exert significant role on the survival, growth, and immune development of neonatal calves [1]. Colostrum contains numerous bioactive constituents, including maternal immunoglobulins, growth factors, antimicrobial peptides, cytokines, and even immune cells, collectively conferring passive immunity against diverse pathogens, promoting intestinal microbiome development, maintaining anti-inflammatory homeostasis, and facilitating immune tolerance to microbiota [1,5-11].

Neonatal calves face notable challenges to their overall health, growth, and survival, particularly due to intestinal inflammation and infectious or non-infectious diarrheal diseases [12,13]. The gastrointestinal tract of neonates is highly susceptible to inflammatory and diarrheal conditions during this critical developmental period [13]. Early colonization of the intestinal tract by microbiota begins in the initial hours and continues through weeks and months of life [14,15]. The diversity of the microbial community is strongly influenced by the inflammatory status of the intestinal environment, and could be further amplified by pathogenic microbiota and their secreted metabolites or factors. Additionally, several evidence suggests that dysregulation of innate immune responses in the gut, coupled with alterations in the intestinal microbiota composition, especially by the colonization of pathogenic flora, significantly contributes to the pathogenesis of inflammatory bowel diseases, leading to persistent diarrhea [15]. Therefore, gaining a comprehensive understanding of the molecular mechanisms underlying colostrum-mediated responses during inflammation and microbiota colonization holds substantial significance in improving calf health, preventing diseases, and developing novel host-directed therapeutic approaches.

Interleukin-6 (IL-6) represents a multifunctional pro-inflammatory cytokine that plays pivotal roles in numerous biological processes, including inflammation, immune regulation, and the interplay between innate and adaptive immune responses. Within the intestinal milieu, IL-6 orchestrates acute inflammation, crucially contributes to immune cell recruitment, and modulates inflammatory processes [16]. Importantly, IL-6 has been identified as a key cytokine involved in the pathogenesis of inflammatory bowel diseases in humans. Various cell types within the intestine, including macrophages, dendritic cells, T cells, and epithelial cells, produce IL-6 in response to diverse stimuli, such as infections, tissue damage, and inflammation [17]. Due to its highly deleterious nature, the production of IL-6 is tightly regulated; however, it can be chronically upregulated under inflammatory conditions, leading to tissue damage, compromised intestinal barrier function, and systemic infections [18].

Recent studies have demonstrated the immunomodulatory activity of COL within the intestinal environment. It has been established that COL can augment the production of anti-inflammatory interleukin-10 (IL-10), thereby favorably shifting immune responses from pro-inflammatory Th1 responses towards anti-inflammatory Th2 responses [19]. Notably, administration of COL in a dextran sulfate-induced colitis model resulted in increased numbers of CD11b-positive leukocytes producing IL-10, ultimately leading to diminished intestinal inflammation via the Src-Akt pathway [20]. Furthermore, IL-10 can negatively regulate IL-6 production by potentially modulating nuclear factor-kappa B (NF-κB) and signal transducer and activator of transcription 3 (STAT3) signaling pathways. However, the precise regulatory mechanisms underlying the interplay between IL-6 and IL-10 in response to colostrum within the intestine remain incompletely characterized.

Our previous studies have elucidated the critical impact of COL administration on the health and development of neonatal Holstein calves [21]. Notably, we observed heightened leukocyte numbers in colostrum-fed calves compared to the control group, providing preliminary evidence of the potential immunomodulatory activity of colostrum [21]. Building upon this, with our interest in comprehensively deciphering the molecular mechanisms underlying COL’s effects in calves, we aimed to employ advanced immunological, cellular biology, and *in vivo* animal models to de-lineate the immunomodulatory properties of COL. In this study, we present the development novel calf-fecal microbiota transplantation (FMT) into convenient C57BL/6 mice models, in which mice harbor engrafted calf microbiota, as an approach to investigate the alterations induced by colostrum within the intestinal environment. Our findings provide experimental evidence regarding the interplay between IL-6 and IL-10 mediated by colostrum in both *in vitro* and *in vivo* models, and propose the utility of colostrum as a preventative or therapeutic tool to mitigate intestinal inflammation in calves.

## 2. Materials and Methods

### 2.1 Ethics

The animal studies and all procedures involved were approved by the State Food and Veterinary Service (SFVS) animal care and welfare committee (approval number: No G2-164) in compliance with the rules of the National Institutes of Health Guide for Care and Use of Laboratory Animals guidelines.

### 2.2 Collection and processing of bovine colostrum

Bovine colostrum was collected from 10 healthy Holstein Friesian cows in their first or second parity from a farm (Lytagra Agriculture company, Naujieji Bernatoniai, Kaunas District, Lithuania). To ensure consistent experimental conditions and eliminate the potential influence of diet or animal management on the results, the colostrum was collected from the same farm where all the animals were managed in a uniform manner, including diet and circadian rhythm. The calves were born within a similar time frame (between 24-48 hours). After the cows gave birth, they were milked every hour for 4 hours to collect early-produced colostrum (10-25 mL). The colostrum samples obtained from different cows were promptly cooled on ice, pooled together, aliquoted into 10 mL portions, and then frozen at −80 °C until the day of the experiments.

#### 2.2.1 Colostrum serum preparation

For cell culture stimulation experiments, the colostrum was prepared following the method described by Zhang et al. with some slight modifications [22]. The pooled colostrum samples were thawed on ice and then centrifugated at 1500 × g at 4 °C for 10 minutes. The resulting pellet, containing cellular components and precipitates, was discarded, and the supernatant was carefully transferred to screw-cap tubes 1.8 mL (JSHD, Jiangsu, China). Subsequently, the tubes were centrifugated at 25,000 × g for 1 hour at 4 °C. The upper fraction of colostrum, which contained fat, was aspirated and discarded, while the middle portion, consisting of colostrum serum, was carefully removed. The colostrum serum was then filtered through a 0.45 µm syringe filter and used for analysis in the experiments.

#### 2.2.2 Colostrum serum protein quantification

The Pierce Coomassie Bradford protein assay kit (Thermo Scientific, #23236) was utilized to quantify and normalize the colostrum samples for the cell culture stimulation experiments. The total protein content in the samples was determined following the instructions provided by the manufacturer. Bovine serum albumin (BSA), which was included in the kit, was used to prepare a standard curve and quantify the protein content of the colostrum samples.

#### 2.2.3 Colostrum inactivation by heat

To generate heat-inactivated colostrum (HI-Col), samples of colostrum serum were subjected to incubation in a water bath at 70 °C for 30 minutes with constant agitation. The HI-Col was subsequently cooled on ice and then centrifuged at 3000 × g for 10 minutes at 4 °C. The resulting supernatant was collected, and the total protein concentration was measured using the Bradford assay Thermo Scientific, #23236). Finally, the HI-Col was utilized for the cell culture experiments.

#### 2.2.4 Ultrafiltration of colostrum

To generate 10, 50, and 100 kDa fractions of colostrum, Amicon-Ultra 15 filters (Sigma-Aldrich, #UFC9010, UFC9050, UFC90100) were filled with colostrum and subjected to centrifugation for 1 hour at 7500 × g, 4 °C. This centrifugation step allowed the separation of colostrum based on molecular size. The flow-through compartment of the filters flows through compartments contained defined molecular size fractions (10, 50, and 100 kDa) of colostrum. The total protein concentration in these fractions was quantified using the Bradford assay Thermo Scientific, #23236). The protein content in each fraction was then used for the cell culture experiments.

### 2.3 Cells and culture conditions

Human Caco-2 intestinal epithelial cells (ATCC HTB-37) were obtained from American Type Culture Collection (ATCC) (https://www.atcc.org/products/htb-37) were cultured in DMEM/F12 supplemented with GlutaMAX (Gibco, #10565018), PenStrep (Gibco, #10378016), and 10 % heat-inactivated FBS (Gibco, #A3160501). Human THP-1 monocytes (ATCC TIB-202) were obtained from the ATCC (https://www.atcc.org/products/tib-202) and maintained in complete RPMI media (Gibco, #11875101) supplemented with GlutaMAX (Gibco, #35050061) and 10 % heat-inactivated FBS (Gibco, #A3160501). Both cell lines were cultured at 37 °C with 5 % CO2. When differentiation of THP-1 cells into macrophages was required, the cells were incubated in complete RPMI media containing 200 ng/mL of Phorbol 12-myristate 13-acetate (PMA) (Sigma-Aldrich, #P8139) for 48 hours. Subsequently, the cells were rested in PMA-free RPMI media for an additional 48 hours before being used in experiments.

### 2.4 Establishment of Caco-2/THP-1 co-culture system

The co-culture experiments were conducted following the methodology described elsewhere with some modifications [23]. Initially, Caco-2 cells were seeded onto permeable 0.4 µm inserts (Thermo Scientific, #141078) and placed in 12-well carrier plates at a density of 2.5 × 10^5^ cells per insert. The cells were then incubated for 12 days, with media changes carried out every two days. The formation of the epithelial layer was confirmed by measuring the transepithelial electric resistance (TEER) using an Evom instrument (NC2120402). The TEER value was considered to be approximately 250-300 Ω.cm2 in DMEM/F12 media, indicating the formation of the epithelium. Subsequently, THP-1 derived macrophages (2.5 × 10^5^ cells/insert) were added to the Caco-2 cell inserts, and the co-culture system was further incubated for 48 hours. During this period, the TEER of the co-culture system was approximately Ω.cm2, indicating the establishment of the Caco-2/THP-1 macrophage co-culture system. All experiments were performed in triplicate.

### 2.5 Stimulation of Caco-2/THP-1 co-culture with colostrum

The Caco-2/THP-1 co-cultures were washed thrice with DPBS (Gibco, #10010023) to remove any residual substances. The colostrum was diluted in DMEM/F12 supplemented with 10 % FBS to achieve a concentration of 250 µg/mL (based on total proteins), and the cells were incubated with the diluted colostrum for 1 hour. After the incubation, PMA (10 µg/mL) was added to the basolateral compartment of the co-culture system, and the cells were further incubated for 24 hours to induce cytokine production. The transepithelial electric resistance (TEER) was measured to assess the integrity of the epithelial barrier. Additionally, the conditioned media from the basolateral compartment was collected and used for cytokine ELISA and LDH assays to analyze the secreted cytokines and assess cell cytotoxicity respectively. In cases where microscopy imaging was necessary, the cells were cultured in 12-well plates without transwell inserts to allow direct visualization. All experiments were performed in triplicate.

### 2.6 LDH release assay

According to the manufacturer’s protocol, LDH activity was measured using the CyQUANT LDH Cytotoxicity Assay Kit (ThermoFisher Scientific, #C20300). In the beginning, 50 µL of conditioned media collected from each sample was utilized for LDH quantification. As a control for maximum LDH release, lysed Caco-2/THP-1 cells were prepared using the supplied lysis buffer. The absorbance of the samples was measured at 490 nm and 680 nm. LDH release was subsequently calculated based on the manufacturer’s instructions.

### 2.7 Immunoblot of NFκB phosphorylation in vitro

The THP-1 derived macrophages or Caco-2/THP-1 co-culture was treated with bovine colostrum and stimulated with PMA as described above. After the treatments, the cells were rinsed three times with ice-cold DPBS (Gibco, #10010023) containing 1 mM sodium orthovanadate (NEB, # P0758S). The membrane from the transwell inserts was cut using forceps and placed in 300 µL of 1X cell lysis buffer (Invitrogen, # FNN0011) for 10 minutes on ice. The membranes were sonicated, and samples were centrifuged at 5,000 × g for 10 minutes at 4 °C. The proteins were quantified using the Pierce BCA protein assay kit (ThermoFisher Scientific, #23227) as described by the manufacturer. The protein samples were incubated at 70 °C in 1X NuPAGE LDS loading dye (Invitrogen, # NP0008) with 1X Bolt Reducing agent (Invitrogen, # B0009) for 10 minutes. The lysates were separated by SDS-PAGE, electrotransferred onto PVDF membrane, and blocked using Intercept TSB blocking buffer (Licor, #927-60001). Membranes were probed with primary rabbit anti-NF-κB p65 (D14E12) (Cell Signaling, #8242) or rabbit-phosphor NF-κB p65 (93H1) (Ser 536) (Cell Signaling, #3033) antibodies, and mouse β-Actin (8H10D10) (Cell Signaling, #3700). Membranes were then probed using secondary IRDye® 800CW Goat anti-Rabbit IgG Secondary Antibody (Licor, #926-32211) for NF-κB or IRDye® 800CW Goat anti-Mouse IgG (Licor, #926-32210) for β-Actin. Blots were visualized using the Licor system.

### 2.8 Fecal microbiome transplantation and generation of calf-FMT C57BL/6 mice

The FMT transplantation (engrafting) and generation of convenient, non-gnotobiotic calf-FMT C57BL/6 mice, containing calf FMT was done as described by Wrzosek et al., with some modification [24].

### 2.9 Depletion of C56BL/6 mice intestinal microbiota

C57BL/6 mice, initially obtained from Jackson Laboratories, are currently maintained in the Animal Facility at the Lithuanian University of Health Sciences (LUHS). Animals were kept according to animal welfare recommendations, 12 h day and night cycle, room temperature at 21 °C and 50 % humidity.

Before the experiments, 8 weeks old, C57BL/6 mice were randomly assigned to different experimental groups, with each group consisting of five mice (n=5). To deplete the bacterial and fungal intestinal microbiota, the mice received penicilin-streptomycin (1g/l) (PEN-STREP, Norbrook, Newry, United Kingdom, #9772) metronidazole (1g/l) (SUPPLIN, 500 mg, SANDOZ pharma, Kundl, Austria #3400), vancomycin (250 mg/l) (Vancosan 1000 mg, MIP pharma, Hamburg Germany, J01XA01) and trimethoprim (500 mg/l) (TRIMETOP 100 mg, Vitabalans Oy, Hämeenlinna, Finland, 3246) in their drinking water for 2 weeks. Additionally, 30 % PEG 4000 (Sigma-Aldrich, Taufkirchen, Germany) solution was administered via oral gavage (200 µl per animal) ant to avoid fungal microbiota fluconazole (12 mg/kg) (Fluconazole, Baxter, Utrecht, Holland) to cleanse the intestine mechanically. During the depletion period, the mice were provided with sterilized food to minimize the introduction of exogenous microbiota. The depletion of endogenous microbiota was monitored by conducting quantitative fecal cultures on Sheep Blood agar plates (Thermo Scientific, #R01200). This allowed the assessing of the reduction in microbial colony-forming units (CFUs) in the fecal samples.

### 2.10 Calve fecal microbiota transplant (FMT) preparation

Fecal material was collected from 7-day-old 3 male and 3 female calves, with a total of 6 calves included in the study. The collected fecal samples were kept refrigerated on the ice during transportation to the laboratory to maintain sample integrity.

Upon arrival at the laboratory, the individual fecal samples obtained from different animals were combined and pooled together. The pooled fecal sample was then diluted in a preservation media (Brain-Heart Infusion broth (Sigma-Aldrich, Taufkirchen, Germany), 20 % skim milk, 1.25 g L-cysteine (Sigma-Aldrich, Taufkirchen, Germany) preservation media medium) to achieve a concentration of 1g/mL of FMT. To ensure homogeneity, the fecal sample was homogenized using the Stomacher 450 Bam homogenization system (Thomas Scientific, #1185V52) for 1 minute at room temperature.

Following homogenization, the fecal microbial transplant (FMT) was divided into aliquots and transferred to sterile polypropylene screw-cap tubes (JSHD, Jiangsu, China). These tubes were then stored at −80 °C until the experiments time to preserve the viability and stability of the microbial content within the FMT samples.

### 2.11 Calve - FMT transplantation to C57BL/6 mice

After the depletion period, the administration of antibiotics through the drinking water was discontinued, and the mice started receiving sterile drinking water to allow for the clearance of antibiotics from their system before the administration of calf FMT.

The necessary number of vials containing calf FMT was defrosted at 10-15 minutes in a water bath at 37 °C and mixed by gently inverting the vials to ensure homogeneity. The calf FMT was then administered to the mice via oral gavage twice a day for 3 consecutive days. Following the last dose of calf FMT administration, the mice were referred to as calf-FMT C57BL/6 mice. These mice then underwent feeding studies or further experiments related to the specific research objectives.

### 2.12 Calf-FMT C57BL/6 mice feeding studies

The calf-FMT C57BL/6 mice were divided into different experimental groups and subjected to feeding with either colostrum (COL) or normal saline (NS), which served as a control.

To administer colostrum, the frozen samples were thawed at room temperature and given to the calf-FMT C57BL/6 mice via oral gavage at a dose of 20 mL/kg. The colostrum was administered every 8 hours twice a day for 5 days.

For the control group, normal saline was administered to the mice at a dose of 20 mL/kg, also via oral gavage. The normal saline was given every 8 hours twice a day for 5 days. After the feeding period, the mice were euthanized by cervical dislocation. Blood samples were collected via cardiac puncture, and the blood was then separated into serum by centrifugation at 10,000 × g for 10 minutes at 4 °C. Furthermore, the intestines of the mice were dissected and used for histological analysis, which involved examining the tissue microscopically to evaluate any changes or effects.

### 2.13 Lethal challenge with S. typhimurium ATCC 14028 strain

The *Salmonella typhimurium* (*S. typhimurium* ATCC 14028) ATCC 14028 strain was cultured on Tryptic-Soy agar (Sigma-Aldrich, Schnelldorf, Germany) plates overnight to obtain single colonies. The bacterial mass was then suspended in sterile saline, and the inoculum size was quantified spectrophotometrically to achieve approximately 1.5 × 108 CFU/mL. Each animal received a volume of 200 µL of the *S. typhimurium* suspension, resulting in an infectious dose of approximately 3.0 × 107 CFU/animal. The accuracy of the inoculum size was confirmed by performing serial dilutions and plating on Tryptic-Soy agar (Sigma-Aldrich, Schnelldorf, Germany) plates (BD Difco, #236920).

24 hours after infection, the calf-FMT C57BL/6 mice received either colostrum (20 mL/kg) or normal saline (20 mL/kg) at regular intervals of 8 hours twice a day for 5 days. In the survival studies, the administration of colostrum or normal saline continued until the experimental endpoint was reached, which was defined as a 20 % loss of body weight or a morbid appearance. The animals were euthanized by cervical dislocation. Blood samples were collected via cardiac puncture, and the blood serum was prepared by centrifugation. The bacterial burden, expressed as log10 CFU/mL, was determined by performing serial dilutions and plating on MacConkey agar plates (BD Difco, #212123)). Additionally, the fecal burden of *S. typhimurium* ATCC 14028 was calculated by serially diluting fecal material collected from the cecum and plating it on XLD agar (Biomerieux, # CM0469B) to differentiate from fecal flora. Sections of the intestine were collected for histological analysis.

### 2.14 Cytokine quantification by ELISA

THP-1 derived macrophages or Caco-2/THP-1 co-cultures were utilized for the cell culture experiments. Following PMA stimulation, the conditioned cell culture media (50-100 µL per assay) collected from the basolateral compartment was subjected to quantification of various cytokines. Specifically, IL-1β (R&D Systems, #DLB50), IL-6 (Invitrogen, #88-7066-88), IL-8 (Invitrogen, #KHC0081), IL-10 (Invitrogen, #EHIL10), IL-12p70 (Invitrogen, #BMS238), TNFα (Invitrogen, #88-7346-88), and INFγ (Invitrogen, #KHC402) were measured using commercially available ELISA kits. As necessary, 96-well plates were coated with capture antibodies, and the assays were performed according to the manufacturer’s instructions.

For the colostrum feeding experiments in calf-FMT C57BL/6, cytokine quantification was performed using 1:5 diluted mouse blood serum. The following cytokines were measured: IL-6 (Invitrogen, #88-7064-88), IL-10 (Invitrogen, #88-7105-88), IL-12p70 (Invitrogen, #88-7121-88), and INFγ (Invitrogen, #88-7314-22). The quantification was carried out following the manufacturer’s instructions.

### 2.15 Evans blue intestinal permeability assay

An Evans blue (EB) solution was prepared by preparing 10 mg/mL of Evans blue (Sigma-Aldrich, #E2129) in saline. The solution was then sterilized by passing it through a 0.4 µm filter to ensure sterility.

For the experimental feeding or *S. typhimurium* ATCC 14028 infection studies involving Calf-FMT C57BL/6 mice receiving colostrum or normal saline, the EB solution was administered to the mice at a dose of 200 mg/kg via intraperitoneal (*i.p*.) route. This administration took place 12 hours prior to the scheduled euthanasia.

To collect samples for EB analysis, the Calf-FMT C57BL/6 mice were euthanized, and their whole intestines were carefully dissected and separated into 4 sections: duodenum, jejunum, ileum and colon. Each section was weighed and homogenized in 500 µL formamide (Sigma-Aldrich, Taufkirchen, Germany, #200-842-0). The EB present in the tissue samples was extracted by incubating the samples at 55 °C for 24 – 48 hours. After incubation, the extracts were clarified by centrifugation at 10000 × g for 10 minutes.

The concentration of EB in the tissue samples was determined spectrophotometrically by measuring the absorbance at 610 nm using a spectrophotometer (Bio-Tek Synergy HT, Santa Clara, USA). The tissue concentration of EB was calculated by using known concentrations of EB in formamide as a reference.

### 2.16 Histological analysis

The calf-FMT C57BL/6 mouse intestines were processed by using Swiss roll technique as described by [25]. Right after euthanasia, the intestines were dissected and placed in ice-cold PBS (Gibco, # 10010031). The cecum was dissected and discarded, and the small intestine was flushed by ice cold PBS, following cold Bouin fixative (5 % acetic acid, 50 % ethanol in deionized water) [21,25]. The intestinal rolls were then formed as described by Grigaleviciute et al., and tissues were placed in tissue processing cassettes, fixed overnight in 10% neutral formaldehyde (Fisher Scientific, # SF98-4), processed and embedded by automatic tissue processor. Tissue sections were then stained with Hematoxylin-Eosin (HE) stain to characterize tissue architecture and Alcian blue (AB) staining to visualize mucins [21]. Tissue sections were then evaluated, and the inflammation as well as tissue architecture was scored by a pathologist using the method described by Erben et. al [26].

### 2.21 Statistical analysis

Data were analyzed using GraphPad Prism (GraphPad Software, San Diego, CA, USA). The statistical significance was analyzed using one-way ANOVA and Mann-Whitney T-tests. Data were considered significant when *p*<0.05.

## 3. Results

### 3.1 Effects of Bovine Colostrum on PMA-Induced Cytokine Production and Caco-2/THP-1 Derived Macrophage Co-Culture Model

To investigate the immunomodulatory effect of COL on cytokine production *in vitro*, we utilized a well-established, polarized Caco-2/THP-1 macrophage co-culture model. The cells were stimulated with phorbol 12-myristate 13-acetate (PMA) that acted as broad-spectrum mitogen inducing various cytokines. Initially, the Caco-2/THP-1 system was pretreated with COL at a fixed concentration of 250 µg/mL for 1 hour. Subsequently, PMA was added at a concentration of 10 µg/mL, in the presence of COL and cell cultures were then incubated for 24 hours to induce cytokine secreation to the culture media withouth noticible phenotype change in cells (Figure 1A).

**Figure 1.**
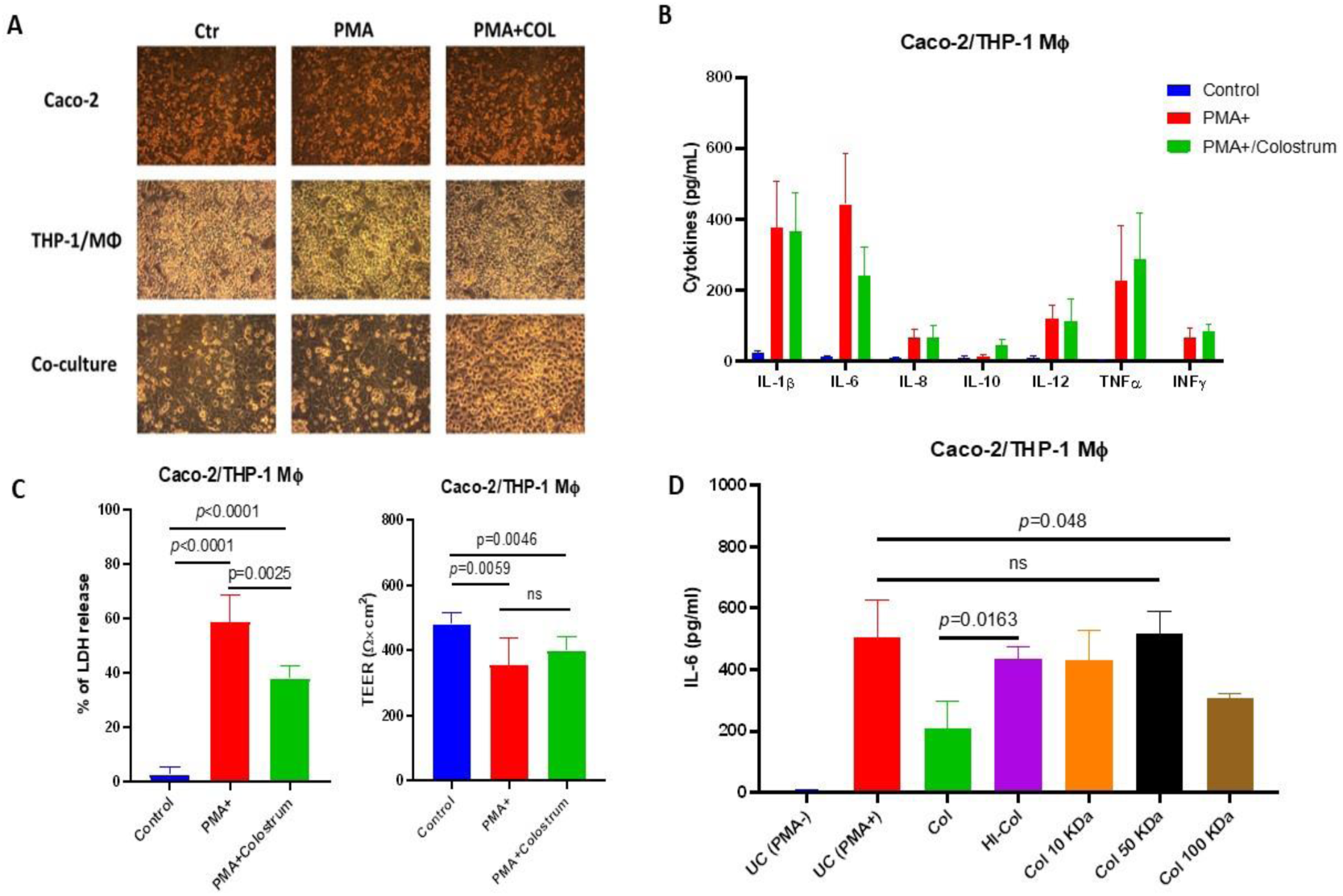
Bovine colostrum exerts immunomodulatory and chemoprotective activity on the Caco-1/THP-1 macrophage co-culture system. Panel A shows the morphological changes of Caco-2, THP-1 and co-cultures during the stimulation with Phorbol 12-myristate 13-acetate (PMA) and treatment with bovine colostrum (COL). Panel B shows the effects of bovine colostrum on PMA-induced cytokine production by Caco-2/THP-1 macrophage co-culture. Panel C shows the effects of bovine colostrum on PMA-induced LDH release and transepithelial electric resistance (TEER) changes. Panel D demonstrates the effect of various treatments of bovine colostrum on the ability to suppress IL-6 production by Caco-2/THP-1 cells.

Our results demonstrated that PMA stimulation significantly increased the production of cytokines, such as IL-1b, IL-6, IL-8, IL-12, TNFα, and INFγ, compared to the unstimulated control (Figure 1B). However, pretreatment of the Caco-2/THP-1 cells with COL resulted in a significant decrease in IL-6 production compared to the cells treated with PMA alone (*p*<0.05). It suggests that COLexerts an inhibitory effect on IL-6 production in response to PMA stimulation (Figure 1B). Furthermore, our study revealed that pretreatment with COL prior to PMA stimulation enhanced the production of IL-10 compared to both the control group and the cells stimulated with PMA alone. These findings indicate that COL has the ability to augment the production of IL-10, an anti-inflammatory cytokine (Figure 1B).

In addition to assessing cytokine production, we evaluated the effects of COL on Caco-2/THP-1 viability by measuring the release of lactate dehydrogenase (LDH) (Figure 1C). Interestingly, we observed that pretreatment with COL prior to PMA stimulation significantly reduced the release of LDH compared to PMA alone. Moreover, PMA stimulation alone or in combination with COL pretreatment reduced the transepithelial electrical resistance (TEER), indicating increased permeability of the Caco-2/THP-1 epithelium (Figure 1C). However, COL pretreatment failed to significantly restore the loss of TEER induced by PMA.

After demonstrating the immunomodulatory activity of COL on IL-6 production using a PMA-stimulated Caco-2/THP-1 cell culture model, we aimed to characterize the types of colostrum constituents responsuble for IL-6 inhibition COL (Figure 1D). To do so, we performed a series of experiments where we used a heat inactivation or ultrafiltration, to prepare a different types of colostrum for the cell culture experiments. Our findings indicated that the IL-6 suppressive effect of COL is influenced by temperature-sensitive factors. Heat inactivation significantly attenuated the colostrum’s ability to inhibit IL-6 production in the Caco-2/THP-1 cell culture model, suggesting the involvement of heat-sensitive components (Figure 1D). To further investigate the factors responsible for the observed IL-6 suppression, we fractionated COL based on molecular size. The fractions with molecular weights of 10 kDa and 50 kDa did not show significant IL-6 inhibitory activity. However, the 100 kDa fraction of COL restored the IL-6 suppressive effect, indicating the presence of immunomodulatory factors with a higher molecular mass (Figure 1D).

These data suggest that COL possess chemoprotective and immunomodulatory activity by suppressing the PMA-induced IL-6 production and cell death by Caco-2/THP-1 derived macrophage model.

### 3.2 Bovine colostrum is able to suppress PMA induced IL-6 production in THP-1 derived macrophages via NF-κB depended and independed manner

After demonstrating the suppressive effect of COL on IL-6 production and its chemoprotective activity, our next objective was to identify wether Caco-2 epithelial cells, or THP-1 derived macrophages are mediating the IL-6 supresory phenotype in response to COL. To investigate this, we individually pre-treated Caco-2 cells and THP-1 macrophages with colostrum and performed co-culture experiments, followed by stimulation with PMA to induce cytokine production and compared the findings to the co-culture experiment.

Interestingly, we observed that COL did not significantly decrease IL-6 production in Caco-2 intestinal epithelial cells alone. However, when THP-1 macrophages treated with COL were stimulated with PMA, a significant reduction in IL-6 was observed. Similar results were obtained when cells in the co-culture system were treated with colostrum and then stimulated with PMA. These findings suggest that the IL-6 suppressive activity of colostrum may be primarily mediated through the THP-1 macrophage axis rather than the Caco-2 epithelial cells.

To further support our observations, we investigated the activation of the NF-κB signalling cascade by measuring the phosphorylation of p65 at Ser536 in THP-1 macrophages and the Caco-2/THP-1 macrophage co-culture after pre-treatment with colostrum followed by PMA stimulation. PMA stimulation, both in monoculture and co-culture conditions, led to increased phosphorylation of p65 at Ser536, indicating NF-κB activation compared to unstimulated controls. However, pre-treatment with colostrum resulted in decreased phosphorylation of p65 at Ser536, specifically in THP-1 macrophages. In contrast, no significant reduction in phosphorylation was observed when THP-1 macrophages were co-cultured with Caco-2 cells, despite the observed decrease in IL-6 protein production in colostrum-treated co-cultures.

These findings provide additional evidence that the IL-6 inhibitory effects of COL are likely mediated through the THP-1 macrophage population, as indicated by the reduction in IL-6 production and decreased phosphorylation of p65 at Ser536. However, the lack of significant reduction in p65 phosphorylation in the co-culture system suggests that other mechanisms may contribute to the observed IL-6 suppression in Caco-2/THP-1 co-cultures treated with colostrum.

### 3.3 Colostrum provides the immunomodulatory activity in the gut and reduces the intestinal permeability

It is widely accepted that the interplay between gut microbiota and the host plays a crucial role in mucosal immune responses and intestinal homeostasis. We first sought to investigate further the histological changes and intestinal inflammation associated with colostrum feeding in calf-FMT transplanted C57BL6 mice. Due to a lack of bovine experimental models and well-established murine models, we conducted experiments using calf fecal microbiota transplantation (FMT) in conventional C57BL/6 mice (calf-FMT C57BL/6).

The C57BL/6 mice were initially treated with a combination of antibiotics and antifungal drugs, followed by PEG 4000 cleansing to deplete the intestinal microbiota. The calf FMT was then engrafted into the conditioned mice via oral gavage. The animals were divided into two groups: one receiving colostrum (COL) (20 mL/kg, q8 hours two times a day) and the other receiving normal saline (20 mL/kg, q8 hours two times a day) for five days. To quantify the feeding-associated changes, we performed experiments using COL and normal saline (NS) feeding and employed the Evans blue assay to assess tissue permeability, which indicates of cellular junction integrity. Our findings revealed that COL significantly (*p*=0.0395) reduced the accumulation of Evans blue as a marker of intestinal permeability in the jejunum and ileum of the calf-FMT C57BL/6 mouse model, as compared to the untreated control (UC) (49.2 and 62.3 μg/g of tissue) or NS feeding (31.2 and 82.1 μg/g of tissue) (Figure 3A). Moreover, feeding with COL was associated with a lower pathological score of inflammation (2 severity score) and noticeable changes in tissue architecture (Figure 3C). Specifically, mice that received colostrum exhibited reduced numbers of tissue-infiltrating neutrophils and macrophages, as compared to those in the UC or NS groups, while mucus production and the localization of Goblet cells were similar (Figure 3B).

**Figure 2.**
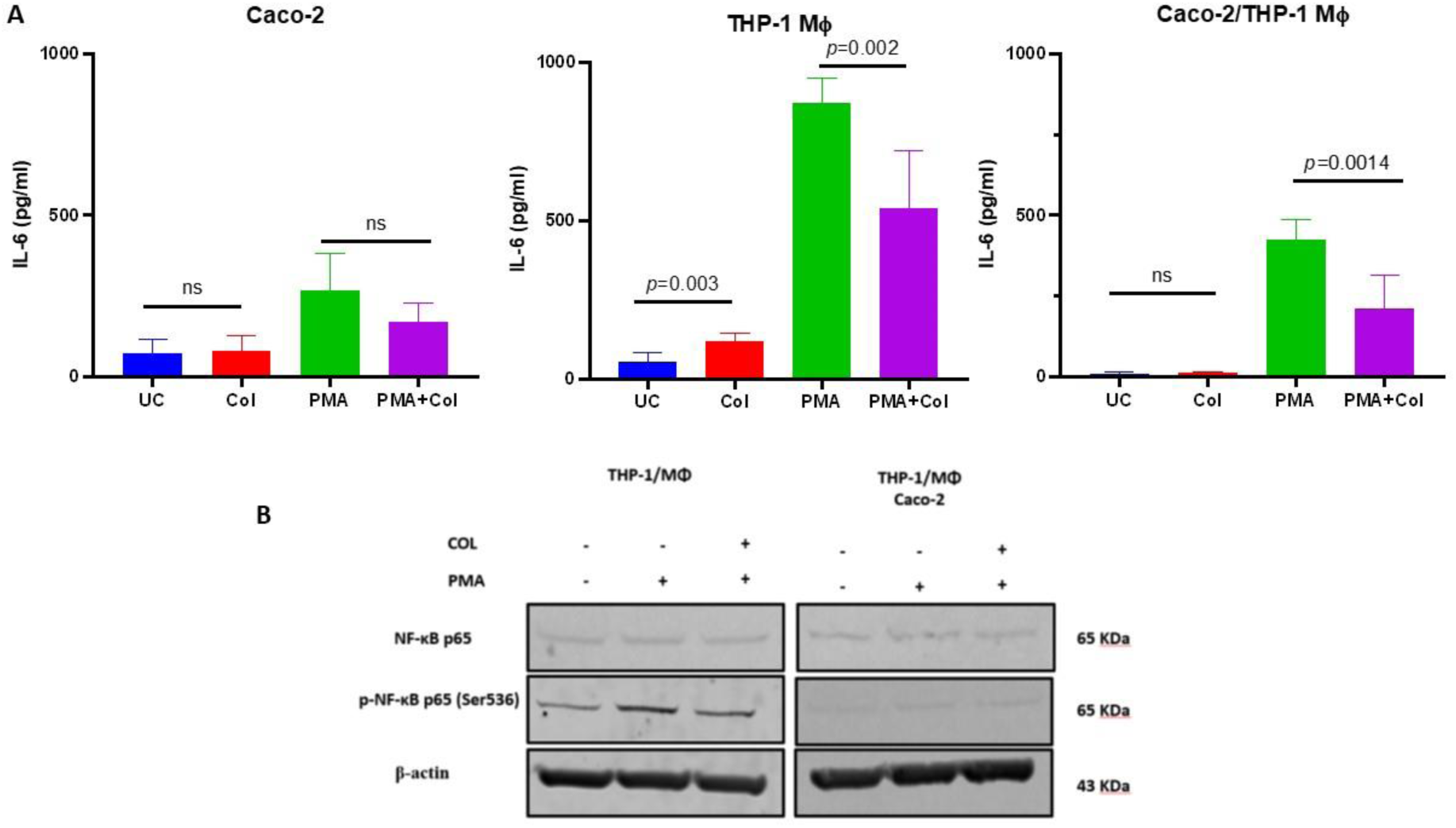
Bovine colostrum is able to reduce IL-6 production via THP-1 derived macrophage modulation. Panel A shows the effects of COL treatment on PMA-induced IL-6 production by Caco-2, THP-1 derived macrophages or their co-cultures. Panel B shows Western blots for NF-κB from THP-1 derived macrophages and Caco-2 co-culture.

**Figure 3.**
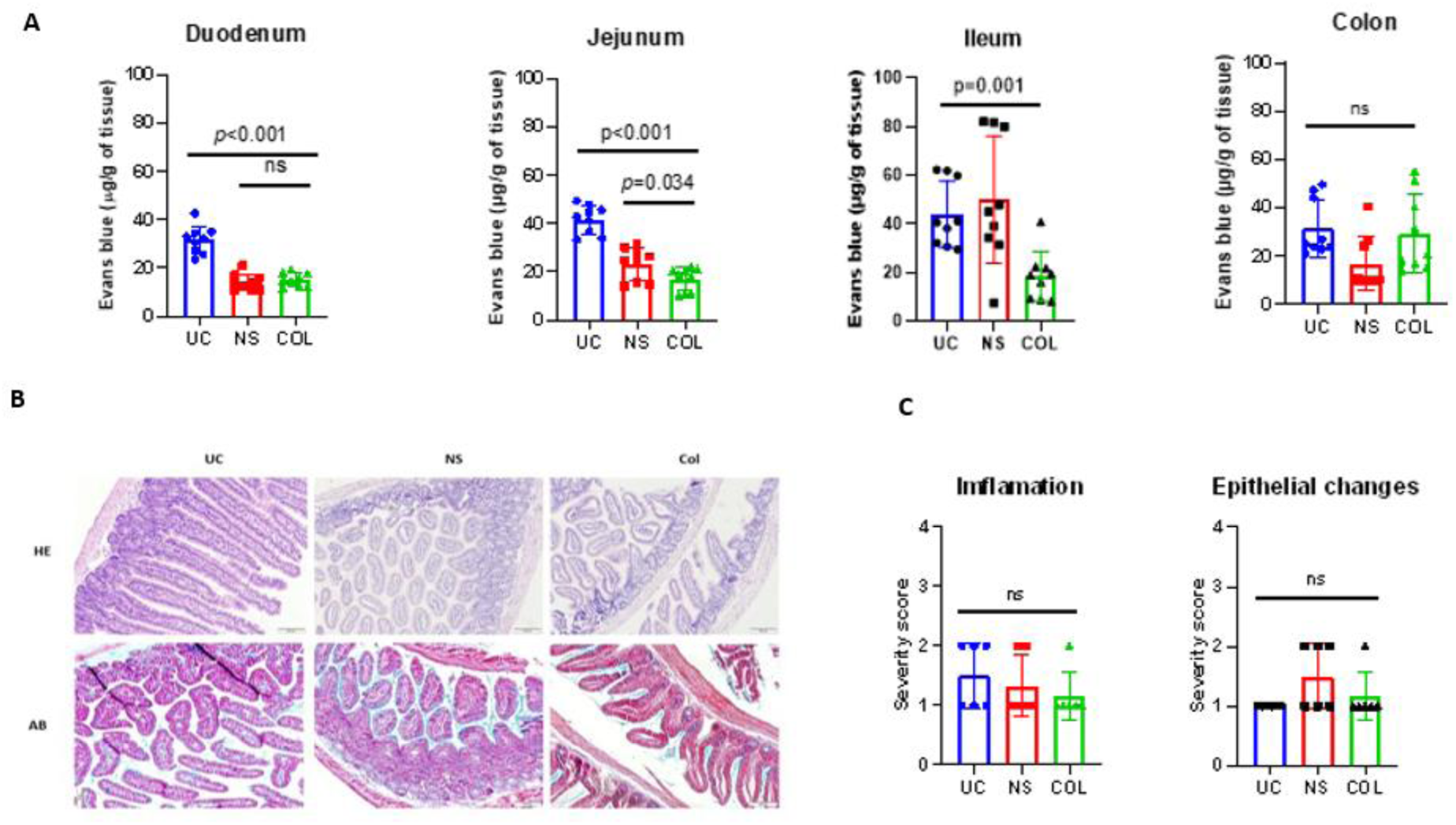
Bovine colostrum decreases the intestinal permeability and infiltration with immune cells in calf-FMT C57BL/6 mice. Panel A demonstrates the intestinal permeability and inflamation measured by Evans blue assay. Panel B shows representative photomicrographs of mouse intestine stained with hematoxilin/eosin (HE) and Alcian blue (AB). Panel C shows the pathological scoring of calf-FMT C57BL6 mice undergoing feeding with normal saline (NS) and colostrum (COL).

Additionally, when the mice were challenged with *S. typhimurium*, a significant (*p*=0.0113) reduction in Evans blue accumulation was observed in the ileum of COL-treated animals in comparison to the UC or NS groups. However, no significant differences were observed in the jejunum, suggesting that the anti-inflammatory and modulatory effects of COL are primarily directed in the ileum. Furthermore, the administration of COL in *S. typhimurium* infected mice resulted in a decreased histological score of inflammation and an enhanced epithelial changes score, suggesting possible pathogen-mediated tissue damage.

Collectivelly, these data suggest that COL modulates the infiltration of immune cells into the gut enviroment as well as reduces the inflammation-mediated damage during lethal *S. typhimurium* challange.

### 3.4 Colostrum exerts immunomodulatory activity in calf FMT-transplanted mice

Surprisingly, the mice fed with COL exhibited significantly decreased production of IL-6 (42.3 pg/ml) compared to the untreated control (UC) or the group receiving normal saline (NS) (Figure 5A). Additionally, we observed an enhanced production of IL-10, indicating that colostrum can modulate the production of cytokines in our calf-FMT C57BL/6 mice (Figure 5A). Building upon this observation, we investigated whether colostrum feeding could modulate inflammatory immune responses in the gut during the challenge with *S. typhimurium* ATCC 14028 (Figure 5B).

**Figure 4.**
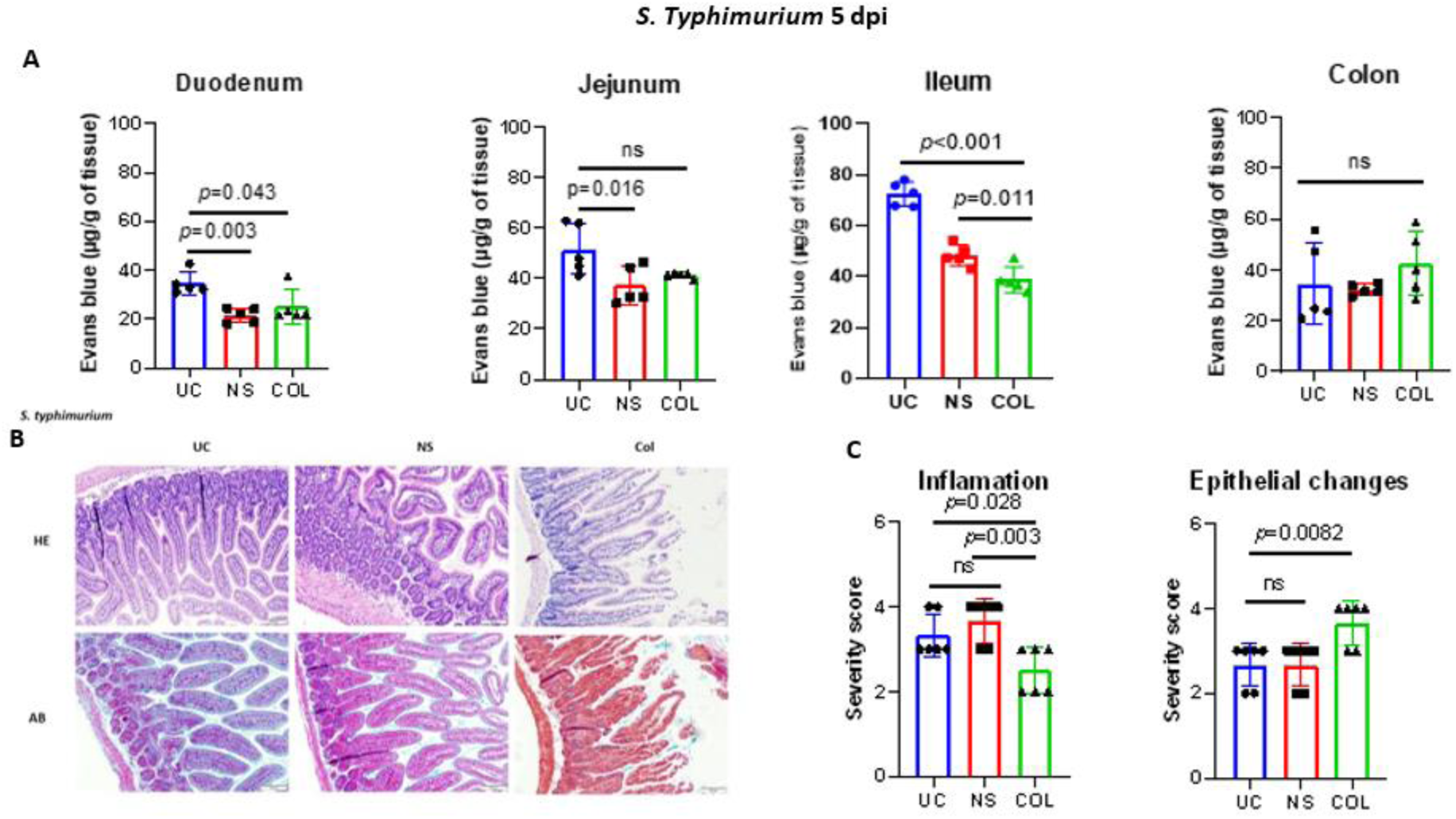
Bovine colostrum reduces the intestinal inflamation in calf-FMT C57BL/6 mice lethally chalanged with *S. typhimurium* ATCC 14028 strain. Panel A demonstrates the intestinal permeability and inflamation measured by Evans blue assay followed lethal chalange with *S. typhimurium* ATCC 14028. Panel B shows representative photomicrographs of *S. typhimurium* infected calf-FMT C57BL/6 mouse intestine stained with hematoxilin/eosin (HE) and Alcian blue (AB). Panel C shows the pathological scoring of *Salmonella* infected calf-FMT C57BL6 mice undergoing feeding with normal saline (NS) and colostrum (COL)

**Figure 5.**
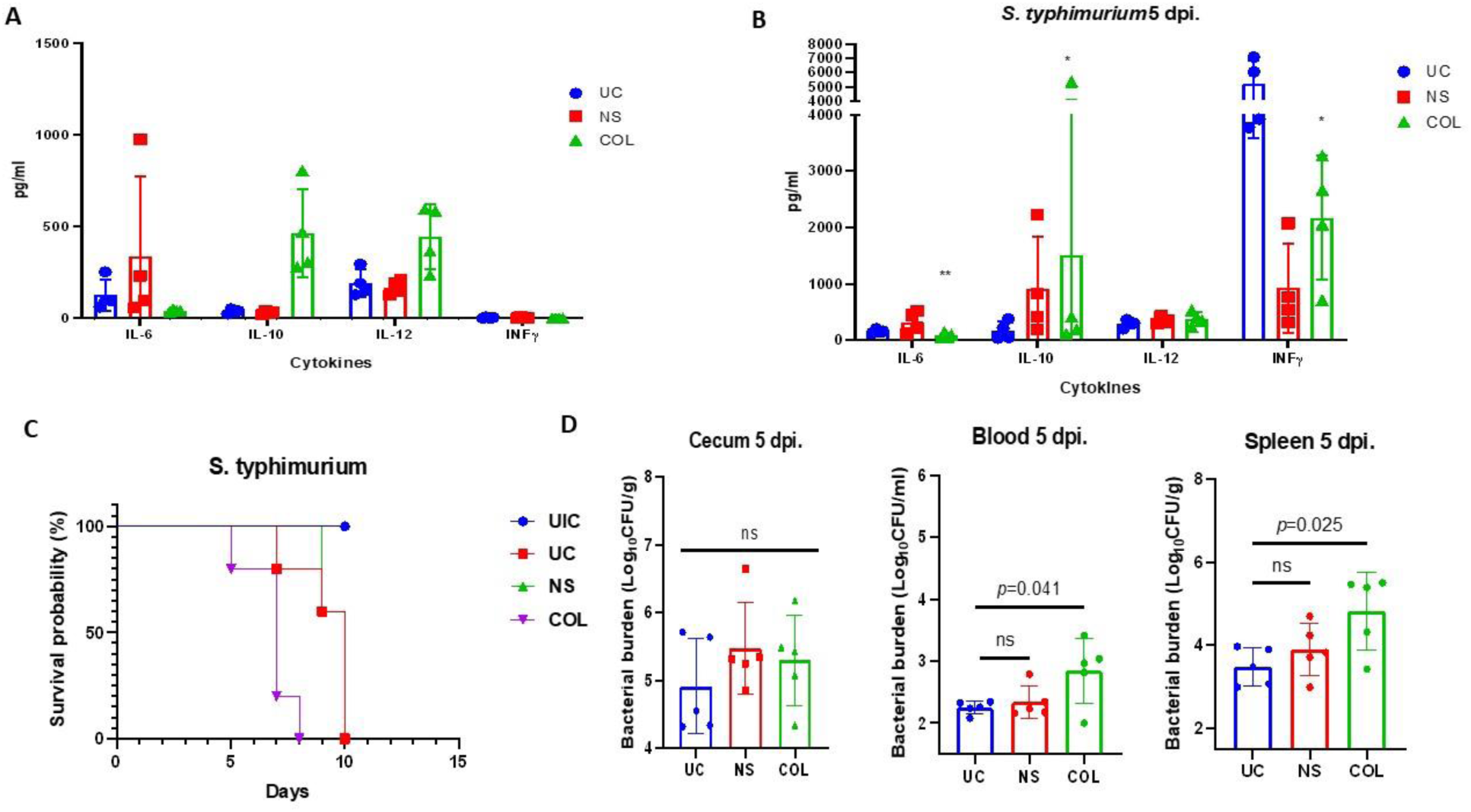
Bovine colostrum exhibits IL-6-targeted anti-inflammatory effects and mitigates *S. typhimurium* ATCC 14028 infection in calf-FMT C57BL/6 mice. Panel A illustrates cytokine production in calf-FMT C57BL/6 mice subjected to colostrum feeding. Panel B displays cytokine production in response to a lethal challenge with the *S. typhimurium* ATCC 14028 strain. Panel C presents survival data of calf-FMT C57BL/6 mice following a lethal challenge with *S. typhimurium* ATCC 14028, with interventions involving normal saline (NS), colostrum (COL), untreated Salmonella-challenged control, or calf-FMT transplanted and non-infected mice. Panel D demonstrates the *Salmonella* bacterial burden subsequent to the lethal challenge and feeding with NS or COL.

To examine this, we infected calf-FMT C57BL/6 mice with *S. typhimurium* ATCC 14028 12 hours before administration of colostrum. The administration of colostrum (COL) (20 mL/kg, q8 hours two times a day) or normal saline (NS) (20 mL/kg, q8 hours two times a day) was continued during the experiment for 5 days. We found that colostrum administration during *S. typhimurium* infection resulted in reduced production of IL-6 and INFγ compared to the untreated control as well as increased production of IL-10 (Figure 5B). However, when a survival study was conducted, the colostrum-treated animals showed decreased survival percentage during the lethal challenge, suggesting an altered intestinal immune defense against *S. typhimurium* and subsequent systemic dissemination (Figure 5C). To confirm this hypothesis, we measured the bacterial burden in the feces (cecum), blood, and spleen (Figure 5D).

Notably, there were no significant changes in cecal *S. typhimurium* burden between infected and treated animals, indicating that COL or NS feeding was not associated with increased bacterial proliferation in the cecum (Figure 5D). Conversely, when assessing systemic dissemination, the bacterial burden in the blood was significantly higher in COL-treated animals (3.4 log_10_ CFU/ml) compared to the UC (2.3 log_10_ CFU/ml) or NS group (2.7 log_10_ CFU/ml). Similar results were observed when quantifying *S. typhimurium* in the spleen of COL treated animals (5.5 log_10_ CFU/ml), suggesting that colostrum feeding provides immunomodulatory activity in the calf-FMT transplanted C57BL/6 mice by modulating IL-6 and IL-10 responses (Figure 5D).

## 4. Discussion

Colostrum (COL), as the initial milk produced by cows after parturition, plays a pivotal role in the systemic and gut immune development of neonate calves, contributing to their enhanced survival and optimal growth during this critical period [27,28]. The rich composition of biologically active substances in COL, including vital nutrients, growth factors, and immunoglobulins, orchestrates the maturation of the intestinal mucosa and shapes the immunological milieu at both local and systemic levels [28]. Notably, COL has been found to influence the microbiome of calves, suggesting a potential interplay between colostrum, the microbiome, and mucosal immune responses that contribute to intestinal homeostasis, tolerance, and protection against diarrheal diseases [29]. Despite the well-documented observations regarding the beneficial effects of COL on calf health and development, there remains a significant knowledge gap concerning the specific molecular pathways regulated by colostrum that underlie its beneficial effects on the intestinal environment and innate immune responses [30–32].

Intestinal inflammation and diarrhea are prevalent and severe conditions in neonate calves, contributing to high morbidity and mortality rates [33]. Various infectious and non-infectious factors can trigger diarrhea, leading to intestinal and systemic in-flammation, compromised intestinal barrier integrity, dysbiosis of the intestinal mi-crobiota, impaired growth patterns, and increased mortality [34,35]. The dysregulated production of pro-inflammatory cytokines in the gut is responsible for initiating and perpetuating the inflammatory processes within the gut leading to systemic inflamation and potential autoimmune disorders [36–38]. A study conducted by Fisher et al. highlighted the critical role of interleukin-6 (IL-6) in bovine diarhetic diseases, suggesting its potential as a prognostic marker [39]. In addition to IL-6, interleukin-1β (IL-1β) and tumor necrosis factor-alpha (TNFα) have been implicated in the acute phase responses observed in neonate calves, underscoring the central role of these cytokines in intestinal inflammation [37].

However, due to the limited avialability of molecular reagents and established bovine cell lines and intestinal models, it is difficult to accurately recapitulate the bovine intestinal environment *in vitro*. Therefore, we employed a well-established Caco-2/THP-1 intestinal model to investigate the innate immune responses to bovine colostrum [40]. To induce inflammatory processes, we utilized phorbol 12-myristate 13-acetate (PMA), a non-specific mitogen that acts through protein kinase C (PKC), nuclear factor kappa-light-chain-enhancer of activated B cells (NF-κB), and mito-gen-activated protein kinase (MAPK) signalling pathways, ultimately resulting in the production of pro-inflammatory cytokines across various cell lines, including Caco-2/THP-1 co-culture [28,41,42].

In our experimental *in vitro* model, pretreatment with COL before PMA stimulation resulted in a decreased production of the pro-inflammatory cytokine IL-6. Interestingly, we observed that COL also exhibited immunomodulatory properties by inducing the production of the anti-inflammatory cytokine IL-10 in response to PMA stimulation, suggesting its potential to regulate the inflammatory response in Caco-2/THP-1 cells. Furthermore, colostrum demonstrated the ability to restore cellular viability by reducing lactate dehydrogenase (LDH) production induced by PMA, indicating its potential cytoprotective effects[43].

Early investigations have provided valuable insights into the potential im-munomodulatory activity of COL. For instance, Biswas et al. demonstrated that exposure of human peripheral blood mononuclear cells (PBMCs) to bovine colostrum led to the polarization of immune responses towards Th1 responses [44]. This suggests that COL has the ability to modulate the balance of immune cell subsets and their associated cytokine profiles. A comprehensive review conducted by Menchetti, highlighted the diverse activities of COL on human innate and adaptive immune responses, indicating its potential therapeutic applications. Colostrum contains high quantities maternal immunoglobulins, cytokines, growth factors, and immunomodulatory milk oligosaccharides [45]. These components have been shown to regulate intestinal mucosal responses through interactions with cytokine and growth factor receptors, as well as with various pathogen-associated molecular patterns (PAMPs) or damage-associated molecular patterns (DAMPs) [46]. This suggests that COL can exert its immunomodulatory effects by influencing the signalling pathways involved in immune cell activation and regulation. The presence of maternal immunoglobulins in colostrum provides passive immunity to newborns, offering protection against various pathogens [9,47,48]. As non protein-derived constituents, immunomodulatory milk oligosaccharides found in colostrum have been shown to interact with the gut microbiota, influencing its composition and function, which in turn can impact immune responses and overall gut health [49].

In our study, we observed that the ability of COL to suppress PMA-induced IL-6 production is influenced by heat, indicating a sensitivity to temperature, which supports the hypothesis of the possible involvement of proteinaceous matter. Furthermore, our subsequent experiments involving ultrafiltration demonstrated that the constituents of COL, capable of passing through 10 kDa molecular cutoff filters, possess the ability to mediate this activity. In contrast to the findings reported by Ann et al., who demonstrated that the inhibition of pro-inflammatory cytokine expression by colostrum occurs through the regulation of epithelial NF-κB, our study reveals that the production of IL-6 induced by PMA in a co-cultured Caco-2 and THP-1 cells probably does not fully rely on the activation of NF-κB [50].

Intestinal inflammation often leads to impaired intestinal barrier function and in-creased gut permeability, which can exacerbate immune responses, disrupt the gut microbiota, and potentially result in systemic infections and mortality, particularly in neonate calves [51,52]. Additionally, colostrum is rich in beneficial probiotic bacteria, such as *Lactobacillus* and *Bifidobacterium*, which have been shown to directly and indirectly impact intestinal health [53,54]. These probiotics can modulate mucus production, maintain tight junctions between epithelial cells, and influence immunological mucosal tolerance through the IL-10/regulatory T cell axis [55]. Thus, the interplay between colostrum and the intestinal microbiota emerges as a critical factor in maintaining intestinal homeostasis.

To validate our *in vitro* findings regarding the IL-6/IL-10 directed immunomodu-latory activity of colostrum, we utilized murine models with engrafted calf fecal mi-crobiota to replicate the intestinal environment of calves in a well-established organism with defined immunological and histological properties. We chose the C57BL/6 background mice due to their wild-type Th1 responses, both at mucosal and systemic levels and similar strategy described by Wrzosek et al. [24]. To simulate acutely inflamed gastrointestinal conditions, we employed a lethal challenge with *Salmonella typhimurium* ATCC 14028 to investigate the effects of colostrum on the host’s innate immune responses to this intestinal pathogen [56]. As *S. typhimurium* induces robust Th1 inflammatory responses in the intestine, characterized by immune cell recruitment, cytokine production, and increased gut permeability, we hypothesized that the immunomodulatory and anti-inflammatory effects of COL would ameliorate the *S. typhimurium* infection, resulting in enhanced lethality due to an inability to mount pro-inflammatory cytokine responses necessary for immune cell recruitment [57].

In our FMT murine model, we observed that COL significantly reduced intestinal permeability in the duodenum, jejunum, and ileum, while having no effect on the colon, as evidenced by reduced Evans blue accumulation in calf-FMT C57BL/6 mice. Surprisingly, we demonstrated decreased intestinal infiltration of neutrophils and macrophages in colostrum-treated animals compared to untreated controls or those receiving normal saline. Following the *S. typhimurium* ATCC 14028 challenge, we observed decreased intestinal inflammation and leakiness in the ileum of colostrum-fed calf-FMT C57BL/6 mice, along with reduced tissue-infiltrating immune cells. These findings suggest the immunomodulatory and anti-inflammatory activity of COL in the calf-FMT C57BL/6 mouse model. Similar insights into the anti-inflammatory properties of colostrum were observed by Menchetti et al., although the exact mechanism was not proposed [45]. Furthermore, we unexpectedly observed that calf-FMT C57BL/6 mice undergoing colostrum feeding exhibited decreased IL-6 production and enhanced IL-10 and IL-12 production compared to untreated calf-FMT C57BL/6 mice or those receiving saline feeding. This finding demonstrates the *in vivo* relevant immunomodulatory activity of COL. When mice were challenged with *S. typhimurium*, colostrum-treated mice displayed an IL-6 suppressive phenotype and increased IL-10 production. This indicates that the antinflammatory activity of COL is directed toward the IL-6/IL-10 axis, even in a highly inflammatory environment, as anticipated, leading to early mortality and systemic bacterial dissemination.

Despite using multiple *in vitro* approaches and validating our findings in novel murine models, our study has several limitations. Firstly, the use of the Caco-2/THP-1 model cannot guarantee that similar phenotypes would be exhibited by bovine epithe-lium. However, due to the lack of established bovine cell lines and intestinal models, as well as limited avialability of immunological reagents, conducting comprehensive *in vitro* studies to evaluate the effects on bovine epithelial models would be challenging. Secondly, calves possess a unique ruminant intestinal system with potentially bovine specific enzymatic and metabolic properties that differ significantly from rodents. Since the intestinal microbiota is known to influence mucosal immune responses, we performed bovine fecal microbiota transplantation (FMT) into mice models with wild-type responses to generate an intestinal environment similar to bovines. Lastly, we understand that our findings in cell culture models and calf-FMT C57BL/6 mice need further validation using bovine models to fully confirm the colostrum-mediated immunomodulatory effects.

## 5. Conclusions

Our study provided an important insight on the immunomodulatory activity of COL in intestinal environment. The findings described in this study suggest that COL has the capacity to regulate the inflammatory response by modulating the balance between IL-6 and IL-10 in cell culture-based models. Furthermore, in our *in vivo* model using calf-FMT C57BL/6 mice, colostrum feeding led to decreased production of IL-6 and enhanced production of IL-10 and IL-12 compared to untreated mice. Moreover, feeding with COL was able to suppress IL-6 production in *S. typhimurium* infected animals, demonstrating powerful immunomodulatory activity during acute inflammation and infection. This confirms the *in vivo* relevance of the immunomodulatory activity of COL and its ability to shift the cytokine profile towards anti-inflammatory responses. Taken together, our data provides valuable insights into the potential therapeutic applications of COL in suppressing inflammation through the IL-6/IL-10 axis. Further studies are warranted to elucidate the underlying molecular mechanisms and validate these findings in bovine models, with the aim of developing targeted interventions for inflammatory disorders.

## Author Contributions

## Institutional Review Board Statement

Conceptualization, R.G. and P.K.; methodology, P.K. and R.G.; software, R.G. and P.K.; validation, R.G. and P.K.; formal analysis, I.P., R.G., P.K.; investigation, R.G., I.P., P.K.; resources, R.G. and P.K.; data curation, I.P., R.G., P.K.; writing—original draft preparation, P.K., R.G.; writing— review and editing, R.P., I.P., E.R.-V., A.B., A.Ž., A.Z., V.Z., A.K., P.M., R.S; visualization, P.K., R.G.; supervision, P.K., P.M. All authors have read and agreed to the published version of the manuscript.

## Informed Consent Statement

Informed consent to collect samples from farm animals were obtained from the representative of the farm. All experiments involving animal subjects were approved State Food and Veterinary Service (SFVS) animal care and welfare committee (approval number: No G2-164).

## Data Availability Statement

All data generated during this study is disclosed within the manuscript. The biological samples obtained during this study are available upon reasonable request from the corresponding author.

## Acknowledgments

In this section, you can acknowledge any support given which is not covered by the author contribution or funding sections. This may include administrative and technical support, or donations in kind (e.g., materials used for experiments).

